# Photothrombotic Ischemic Thalamic Stroke in Mice Recapitulates Spontaneous Pain Features of Central Post-Stroke Pain in Humans

**DOI:** 10.1101/2025.06.26.661797

**Authors:** Jeremy B. Ford, Olive Tambou Nzoutchoum, Jarret A. Weinrich, Debleena Chatterjee, Allan I. Basbaum, Jeanne T. Paz

## Abstract

Central post-stroke pain (CPSP) is a highly distressing condition that develops in 50% of people who suffer a thalamic stroke, and is typically unresponsive to current clinical treatments. Hypoxic damage to the ventral posterolateral (VPL) and ventral posteromedial (VPM) sensory thalamic nuclei, in particular, precipitates CPSP. One barrier to developing treatments for CPSP is the lack of preclinical models of thalamic ischemic stroke. In this study, we present a novel mouse model of CPSP induced through targeted photothrombotic ischemia. After eliciting hypoxia in the sensory thalamus of male mice, we assessed pain behaviors over a four-week period. Stroke-affected mice exhibited a persistent spontaneous facial grimace from day four to week four post-stroke, indicative of pain. Hind-paw mechanical hypersensitivity indicative of altered nociception, characteristic of VPL and VPM hemorrhagic CPSP models, was not detected in our model. Immunofluorescence analysis revealed increased activated microglia (Iba1) and reactive astrocytes (GFAP). Iba1 fluorescence intensity in the VPL thalamus–but not the VPM thalamus–correlated with the severity of facial grimace at four weeks post-stroke. Clustering based on behavioral phenotypes identified a subpopulation of mice in which grimace pain spontaneously resolved, by four weeks post-stroke, relative to sham controls, suggesting that this model can be used to understand how stroke recovery may influence pain chronification. This model provides a valuable tool to investigate the cellular and circuit mechanisms underlying CPSP after an ischemic thalamic stroke.

**Significance Statement:** Research into central post-stroke pain (CPSP) is hindered by the lack of preclinical models that match the clinical presentation of the condition. Although 87% of clinical strokes are ischemic, current CPSP preclinical models induce hemorrhagic strokes. To our knowledge, our mouse model is the first to induce CPSP by photothrombosis in the ventral posterolateral and posteromedial thalamus. Moreover, our study quantifies spontaneous pain, using a facial grimace score, in addition to nociception, whereas previous CPSP studies have relied exclusively on nociceptive tests. Finally, our model recapitulates some of the variability observed in clinical CPSP. We anticipate that this model will help dissect the roots of chronic pain after a thalamic stroke, and facilitate development of novel therapies for CPSP.

## Introduction

There are over 12.2 million new strokes globally each year (Feigin et al. 2022). Since both the total incidence of stroke and the number of years with lived disability are increasing (Tsao et al. 2023), the number of stroke survivors that exhibit secondary sensory, motor, and cognitive issues is growing. Pain is extremely common following stroke, occurring in up to 70% of patients (Ali et al. 2023); up to 35% of patients (Betancur et al. 2021) will develop central post-stroke pain (CPSP), described as “among the most spectacular, distressing, and intractable of pain syndromes” (Henry, Lalloo, and Yashpal 2008). These patients develop severe squeezing, burning, and stabbing sensations (Urits et al. 2020) in the months following stroke (G. Andersen et al. 1995), which can last for decades (Tamasauskas et al. 2025). Nearly 50% of thalamic strokes result in CPSP, most commonly due to strokes in the ventral posterolateral (VPL) and ventral posteromedial (VPM) nuclei (Treister et al. 2017; Liampas et al. 2020).

To date, there are no effective therapies that reliably mitigate the pain in CPSP patients, or prevent its onset. Medications that are typically used for other chronic neuropathic pain states, including complex regional pain syndrome (Treister et al. 2017) and spinal cord injury (Teasell et al. 2010), are prescribed for CPSP patients, with first line treatments involving anti-seizure medications (e.g. pregabalin) and antidepressants (e.g. amitriptyline), but these have demonstrated mixed effectiveness (Liampas et al. 2020; Ri 2022; Treister et al. 2017). Progress in the development of better pain management in CPSP patients has been hampered by limited preclinical models that adequately reflect the clinical presentation of the condition.

Current preclinical models of CPSP induce a hemorrhagic stroke in the VPM or VPL thalami using intracerebral injection of collagenase IV, which break downs the extracellular matrix and the basal lamina of blood vessels (Gritsch et al. 2016; MacLellan et al. 2008). However, 87% of strokes in patients are ischemic (Tsao et al. 2023), and ischemic and hemorrhagic strokes differ in both the clinical presentation and the timeline of recovery (K. K. Andersen et al. 2009; Chae, Zorowitz, and Johnston 1996). In addition, to date, preclinical studies monitored only stimulus-provoked measures of pain (e.g. mechanical hypersensitivity/allodynia), which only occurs in ~60% of CPSP patients (Ri 2022). These studies did not assess spontaneous/ongoing pain, which is experienced by up to 85% of patients (Klit, Finnerup, and Jensen 2009). Here, we present a novel mouse model of CPSP induced by photothrombotic ischemic stroke in the VPL and VPM thalamic nuclei, and demonstrate significant, long-lasting spontaneous pain using the facial grimace scale.

## Materials and Methods

### Experimental Design

#### Animals

All animal protocols were approved by the Institutional Animal Care and Use Committee at the University of California, San Francisco and Gladstone Institutes. Precautions were taken to minimize stress as well as the number of animals studied. Adult, C57Bl/6 male mice were purchased from Jackson Labs (Stock No. 000664). Mice were group-housed with same-sex littermates in cages and fed ad libitum. Animals were kept in a 12-hour light/dark cycle (lights on 7:00 AM to 7:00 PM) and mice weighed 25.3 ± 1.2 g before stroke or sham surgery.

We first established optical parameters to induce photothrombotic stroke (described below), in 25 mice (12 shams and 13 strokes). Following protocol optimization, pain behavior testing and histological characterization of the stroke size, location, and neuroinflammation was performed in 30 mice (n = 13 shams and 17 strokes) across three replicate cohorts.

#### Thalamic photothrombotic stroke

Mice were weighed and anesthetized with 4%–5% isoflurane in oxygen, after which they were placed in a stereotaxic frame with the skull leveled between bregma and lambda. While maintained at 3% isoflurane, we drilled a 0.5 mm burr hole in the skull, through which a 0.2 mm diameter optical cannula (CFML12U-20, Thorlabs Inc.), connected to an optical fiber, was slowly (~25 m per second) lowered into the thalamus using the following coordinates: −1.54 mm from bregma, +1.84 mm lateral from midline, and 3.0 mm below the cortical surface (target area, VPL and VPM thalamus) (**Fig. 1**). Next, mice received an intraperitoneal injection of Rose Bengal dye (RB, 40 mg/kg) (Item 330000-5G, Sigma-Aldrich, Millipore Sigma), diluted in sterile saline (8.0 mg/mL) (NC9054335, Fisher Scientific, Thermo Fisher Scientific Inc.), as described in our previous studies (Paz et al. 2010; Necula et al. 2021; Klein et al. 2025; F. S. Cho et al. 2022). To induce focal ischemic stroke, a 532 nm laser (MGL-III-532-200mW, Opto Engine LLC) coupled to the optical fiber was used to irradiate the brain at 3 mW, delivering optical doses of 95 J/mm^2^ (16 min, 35 sec), 120 J/mm^2^ (21 min), or 190 J/mm^2^ (33 min). Sham mice had the optical fiber placed in the VPL and VPM thalamic nuclei, and received laser irradiation with an equivalent volume of saline instead of RB. All strokes were induced in the right hemisphere. Post-operative pain was managed with sustained-release buprenorphine (3.25 mg/kg, subcutaneously; Lot #s B230045, B230102, B220119, B210109A, B210108, AB210078; Ethiqa XR, Fidelis Animal Health). Sustained-release buprenorphine, which provides analgesia for up to 72 hours, was administered to mitigate surgery-induced pain.

**Figure 1.**
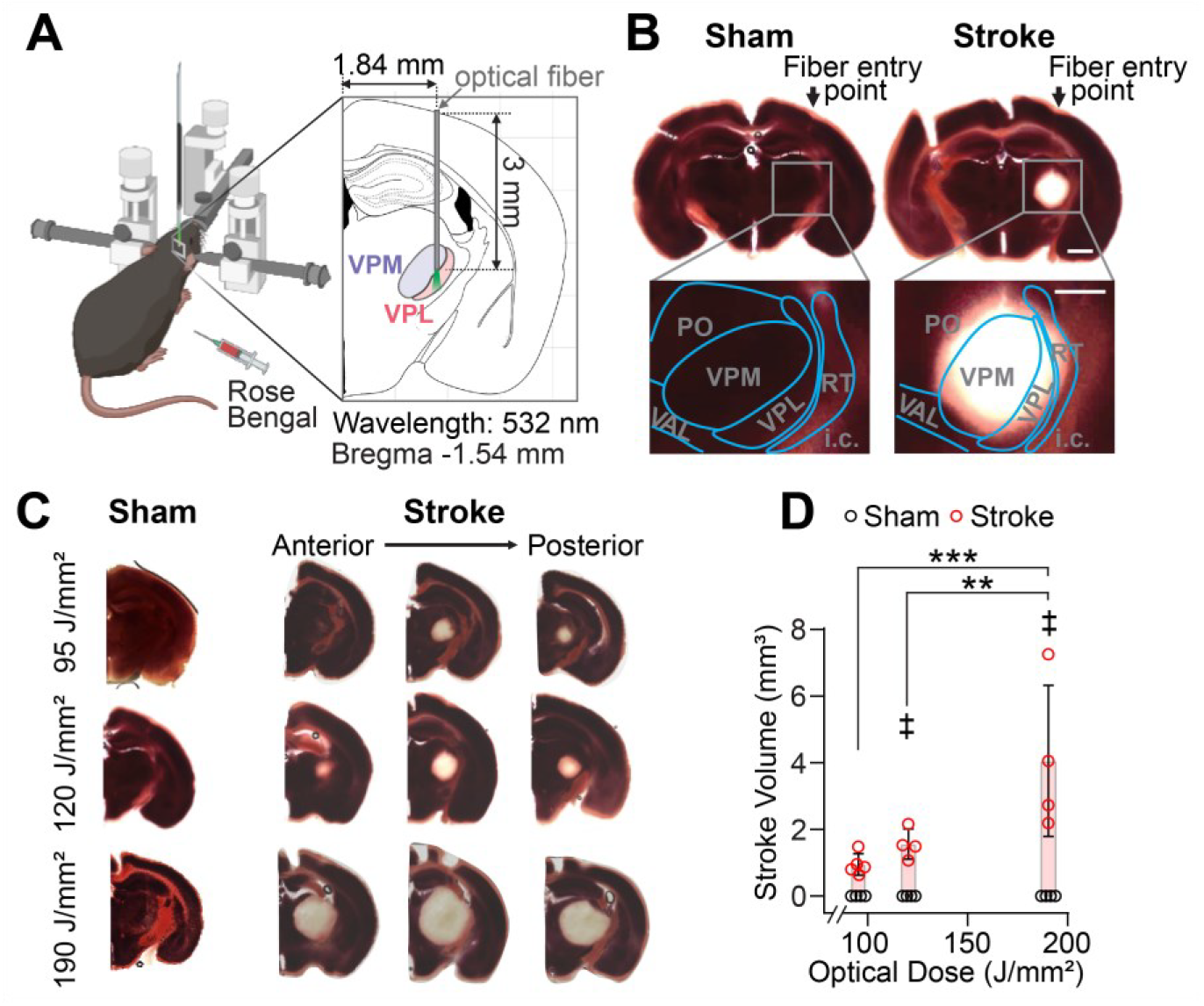
Validating hypoxia in the thalamus. **A)** Schematic of photothrombotic stroke induction. An optical fiber is placed in the right VPL and VPM thalamus under stereotaxic guidance. Either Rose Bengal (stroke group) or an equivalent volume of saline (sham group) was injected (i.p.) prior to optical irradiation at various light doses. **B)** Coronal brain sections stained with TTC 24 hours after irradiation. TTC staining fails to turn red in hypoxic areas, demonstrating stroke induction in the VPL and VPM thalamus, but not in sham mice. Insets show higher magnification on the thalamic regions to illustrate the stroke location. **C)** Representative coronal brain sections from mice exposed to different optical doses, stained with TTC 24 hours after irradiation. Multiple sections from each stroke mouse are included to provide a view of the extent of hypoxia. **D)** Stroke volume follows a dose dependence where greater light doses lead to larger stroke volume. Data represent all points from min to max from n = four to five sham (black circles) or stroke (red circles) mice per optical dose. Two-Way ANOVA with Tukey’s Multiple comparisons. **p < 0.01, ***p<0.001, ‡ p < 0.05 between stroke and sham at the same dose.

#### Behavioral Test Design

Mice had either stroke or sham procedures performed with an optical dose of 120 J/mm^2^. Mice were tested at baseline, four days, and one, two, and four weeks after stroke using the facial grimace scale (Langford et al. 2010), open field test (H. Cho et al. 2013), von Frey test (Chaplan et al. 1994), and rotarod test (see Extended Data). Mice acclimated to the testing room for at least 45 minutes prior to running any tests. At least one hour of rest was provided between behavioral tests, which were administered in order from least to most noxious: the facial grimace scale, followed by the open field test, and then the von Frey test. Rotarod testing was performed on a separate day at baseline, one week, and four weeks post-stroke.

#### Facial Grimace Score

We used the grimace score to assess spontaneous pain (Langford et al. 2010). Mice were placed in a small enclosure (4” x 4” x 5.5”) (Item # 57824, Ugo Basile, Stoelting Co.) with a transparent wall for 5 minutes and were recorded with a camera (Item 960-001105, Logitech) at 60 frames per second. Videos were manually scored, on a scale of zero to two, by selecting frames with clear views of multiple angles of the face, and applying the facial grimace metrics to identify grimace features in the eyes, nose, cheeks, ears, and whiskers (**Fig. 2A, Fig. 2-1A–F**). A score of zero indicated no grimace feature; one indicated a mild feature (present but subtle or only slightly deviating from normal); and two indicated a strong feature (clearly visible with a large deviation from normal). To qualify as a true grimace, identified features had to be present throughout the entire video. A single grimace score for each mouse was calculated by summing the scores of the five metrics.

**Figure 2.**
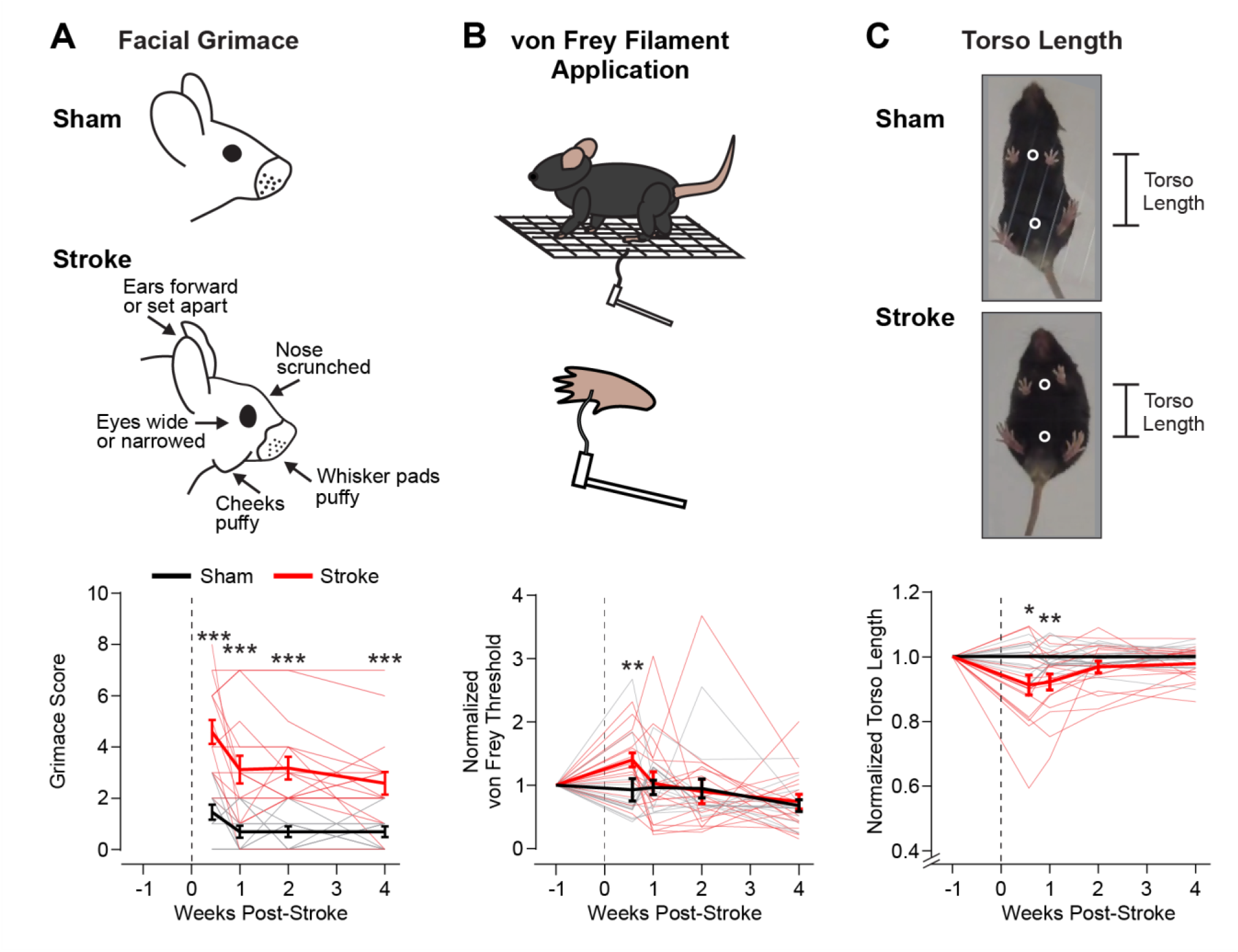
Ischemic thalamic stroke induces acute and chronic pain behaviors in mice. **A)** Facial grimace score. *Top:* The facial grimace is measured by summing scores assigned to the eyes, ears, nose, cheeks, and whiskers. *Bottom:* Evolution of the facial grimace score from four days to four weeks post-stroke in stroke and sham mice. **B)** von Frey Filament assay. *Top:* The assay to the applies von Frey filaments plantar surface of the left (contralateral) hind-paw to calculate the mechanical pressure threshold to elicit a nocifensive response. *Bottom:* Evolution of the von Frey threshold after stroke, normalized for each mouse to its pre-stroke threshold. **C)** Torso length. *Top:* Torso length, a indicative of hunched posture exhibited when mice are in pain, is measured while mice move freely in an open field. *Bottom:* Evolution of torso length, normalized to baseline for each mouse, and normalized to shams within each cohort. Thin traces represent individual mice. Thick traces represent mean ± SEM for each group (stroke: red, sham: black). Data show all points from min to max. Repeated measures Two-Way ANOVA (α = 0.05) with Tukey’s multiple comparisons (*p < 0.05, **p < 0.01, ***p < 0.0001).

#### Mechanical nociception test

Mice were tested for sensitivity to mechanical stimulation of the hind-paws (Shields, Eckert, and Basbaum 2003) using von Frey filaments (Item # 58011, Stoelting Co.). Mice were placed into a multi chamber setup (Item # 57824, Ugo Basile, Stoelting Co.) on top of a gridded mesh platform (Item # 57816, Stoelting Co.) which provided access to the left hind paw (**Fig. 2B**). Starting with the 0.6 g filament, filaments were applied for ~five seconds using the Dixon up-down method (Dixon 1980; Chaplan et al. 1994) to identify the pressure threshold for a nocifensive response, defined as a rapid paw withdrawal or licking, shaking, or stamping of the paw at the onset or offset of the filament. At least one minute separated each filament application. This procedure was repeated twice, yielding three total trials per test day. The pressure resulting in a 50% chance of a nocifensive response was calculated using the established equation (Chaplan et al. 1994) and then averaged across the three trials. The von Frey test was conducted at least two separate times prior to stroke, and again at four days, as well as one, two, and four weeks after stroke. To establish a baseline value, nocifensive response thresholds were averaged across the pre-stroke time points. Results were normalized to baseline to monitor individual responses in each mouse.

#### Open Field test

Mice were placed in a cylindrical chamber (diameter = 30 cm, height = 40 cm) with a transparent acrylic floor mounted above a camera. Mice were allowed to move freely for 10 minutes while the camera recorded from below at 48 frames per second. Videos were analyzed using a custom pipeline incorporating Deeplabcut (Mathis et al. 2018), as described below, to track mouse location and specific anatomical features.

Open field video analysis used a combination of automated anatomical feature tracking with Deeplabcut and calculations performed in MATLAB (MATLAB Version 24.1.0.2653294 (R2024a) Update 5, Mathworks). For the Deeplabcut model, a set of ten videos, spread across time points, with equal number of stroke and sham videos, were selected, and cropped so that only the cylinder and immediate surroundings were visible. A total of five points along the midline were manually annotated in each extracted frame. These were: Nose, Chest, Mid-Trunk, Abdomen, and Tail Base. The model was first trained on 30 frames per video, over 400,000 iterations, and this was refined twice, increasing to 60 frames per video, with 1,000,000 total iterations. This model was then used to determine the locations of the listed anatomical points for each frame across all videos.

Tracked anatomical points for each frame were trimmed so that only the first ten minutes were analyzed. Basic metrics relating to distance traveled (activity and mobility readout), time in cylinder center (anxiety readout), and maximum velocity (mobility readout) (Seibenhener and Wooten 2015) were tracked from the open field test (described in Extended Data) (**Fig. 2-2A–I**). Bouts of motion were identified based on: 1) a velocity threshold, defined by the average velocity of the Chest, Mid-Trunk, Abdomen, and Tail Base points, and 2) a minimum duration of one second. Frames meeting these criteria were used to calculate the distance between the chest and abdomen points (**Fig. 2C**), and these values were subsequently averaged. Torso length was assessed exclusively during bouts of motion to avoid confounding effects of sitting or rearing. Because mice innately have different body lengths, all torso length measures were normalized to each mouse’s pre-stroke baseline length. In one cohort, it was identified that the resolution of the camera was reduced at the pre-stroke recording, resulting in an increase in the baseline normalized torso length at post-surgical time points (**Fig. 2-1G–I**). This artificially increased the effect size of torso length differences between stroke and sham mice at four weeks post-stroke. This was overcome by normalizing the torso length of all mice to the average value of torso length for sham mice within the same cohort at each time point.

### Histology

#### 2,3,5-Triphenyltetrazolium chloride (TTC) staining

Twenty-four hours after stroke inductions, mice were transcardially perfused with cold 1x phosphate buffered saline (PBS) (Ref 46-013-CM, Corning Inc.), and brains were excised and sectioned in cold PBS using a vibratome to produce 0.5 mm thick sections. Sections were immediately placed into a prepared 2% (w/v) 2,3,5-Triphenyl-tetrazolium chloride (TTC) (Item T8877-10G, Sigma-Aldrich, Millipore Sigma) solution in PBS, kept at 37^°^C, and incubated for 10 minutes while covered with foil to block light exposure. Following incubation, sections were washed in PBS, then transferred to 4% paraformaldehyde (PFA) (Catalog # J19943-K2, Lot # 242168, Thermo Fisher Scientific Inc.) for one hour, then 30% sucrose for at least one hour. Last, sections were mounted on a microscope slide and brightfield imaging was performed using a widefield microscope (BZX710, Keyence Corporation of America). Using ImageJ, the stroke area of each section was calculated. Stroke volume was then calculated by summing the infarct areas over all sections and multiplying by the slice thickness (Area of infarct (mm^2^) x slice thickness (mm)) (Swanson et al. 1990; Hawkins et al. 2014; Tian et al. 2023).

#### Immunohistochemistry

Mice were euthanized with sodium pentobarbital (39 mg/mL) (Fatal Plus, Vortech NDC 0298-9373-68, Lot # 2947) following completion of all behavioral tests (n = 7 shams and 12 stroke at seven-weeks post-stroke; n = 6 shams and 5 stroke at 22-weeks post-stroke). Brains were excised following cardiac perfusion and fixed in 4% PFA (Lot # 242168) overnight and stored in 30% sucrose. Tissue was cut into 25 um coronal sections using a microtome (SM2010R, Leica Inc.). Sections were stained for markers of inflammation, using a two-stage staining protocol with primary antibodies for reactive astrocytes (1:1000 chicken anti-GFAP, ab4674, abcam) and activated microglia (1:500 rabbit anti-Iba1, 019-19740, Fujifilm Wako Pure Chemical Corporation), and secondary antibodies (1:1000 Alexa Fluor 488 goat anti-chicken, A-11039, Invitrogen, Thermo Fisher Scientific and 1:500 Alexa Fluor 555 goat anti-rabbit, A-21428, Invitrogen, Thermo Fisher Scientific). Sections were mounted on slides in VectaShield mounting media with DAPI (H-1200-10, Vector Laboratories, Inc.), and then imaged on a widefield fluorescence microscope with using 2x and 10x objectives (BZX710, Keyence Corporation of America).

#### Image segmentation

For bulk histological analysis, fluorescence images were imported into FIJI (Schindelin et al. 2012). Sections containing the center of the stroke core, which was observed as high Iba1 staining surrounded by high intensity GFAP staining (**Fig. 3A–B**), were used for analysis. Sections containing inflammation from optical fiber placement were chosen for sham mice. The thalamus ipsilateral (right) and contralateral (left) to optical dose delivery were each manually segmented. The region of interest (ROI) corresponding to the stroke core in each section was manually segmented based on the outer edge of Iba1 staining and the inner edge of the glial scar labeled with GFAP. This was then used to estimate the cross-sectional area of the stroke core, and to understand how each thalamic nucleus was affected (described below). The posteriority, laterality, and depth of the stroke core was estimated as the Allen Brain atlas anterior-posterior coordinate of the brain section, and the midline laterality and depth from cortical surface of the centroid of the stroke core, respectively. The lateral ventricle in each hemisphere was manually segmented, and the cross-sectional areas were used to calculate the ipsilateral/contralateral (Right/Left) ventricle area ratio for each section.

**Figure 3.**
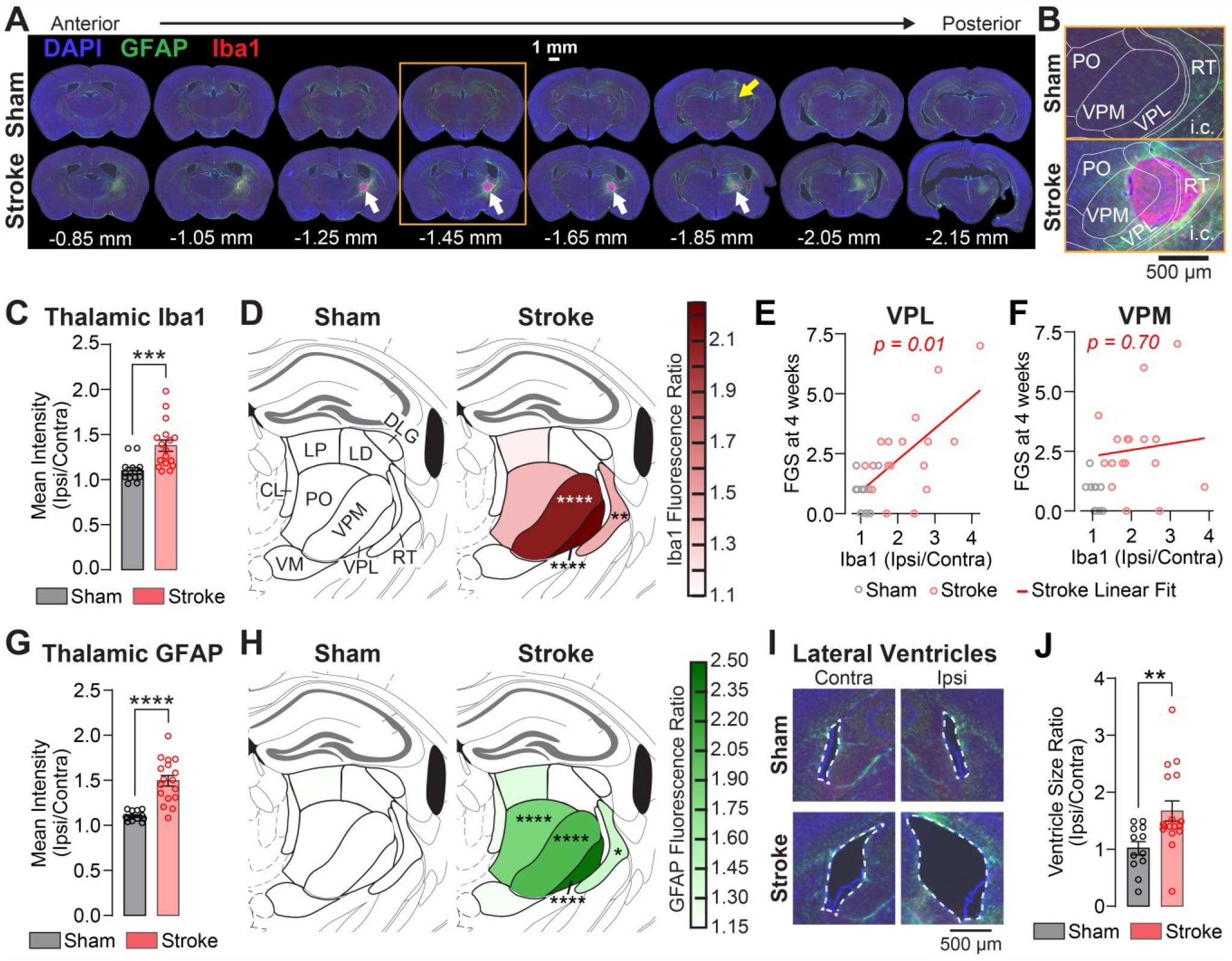
Thalamic photothrombotic stroke elicits neuroinflammation and brain morphological changes. **A)** Representative staining of reactive astrocytes (green), microglia (red), and nuclei (blue) from a sham and a stroke mouse across the thalamus. Stroke core (white arrows) and inflammation due to optical fiber insertion (yellow arrow) are highlighted. **B)** Overlay of the Allen Brain Atlas on the sensory thalamus in sections from (A) outlined in the orange box. **C)** Mean fluorescence intensity of Iba1 within the thalamus in sections containing optical fiber placement (sham mice) or the stroke center (stroke mice), normalized to the mean fluorescence intensity in the contralateral thalamus. **D)** Mouse brain atlas showing the distribution of Iba1 staining quantified over different thalamic nuclei for sham (left) and stroke (right) mice according to the shown color map. Color values represent the ipsilateral/contralateral fluorescence ratio for each thalamic nucleus. **E)** Scatterplot of the facial grimace score at four weeks post-stroke versus the Iba1 fluorescence intensity (ipsilateral/contralateral ratio) in the VPL thalamus for sham (gray circles) and stroke (red circles) mice. A linear regression best fit line with p-value is shown for stroke mice, whereas sham mice did not demonstrate a significant linear relationship. **F)** Scatterplot of the facial grimace score at four weeks post-stroke versus the Iba1 fluorescence intensity (ipsilateral/contralateral ratio) in the VPM thalamus for sham (gray circles) and stroke (red circles) mice. A linear regression best fit line with the associated non-significant p-value is shown for stroke mice to compare with the regression in the VPL (**E**). The non-significant regression for sham mice is not shown. **G)** Mean fluorescence intensity of GFAP within the thalamus in sections containing optical fiber placement (sham mice) or the stroke center (stroke mice), normalized to the fluorescence signal in the contralateral thalamus. **H)** Mouse brain atlas showing the distribution of GFAP staining quantified over different thalamic nuclei for sham (left) and stroke (right) mice according to the shown color map. Color values represent the ipsilateral/contralateral fluorescence ratio for each thalamic nucleus. **I)** Representative cross-sections of the left and right ventricles of a sham and a stroke mouse. **J)** The cross-sectional area of the ipsilateral lateral ventricle, normalized to the cross-sectional area of the contralateral lateral ventricle, in sham and stroke mice. Data represent all points from min to max, Mean ± SEM, with a two-sample t–test (**C**,**G**,**J**), or a Tukey’s multiple comparisons following a positive Two-Way ANOVA (**D**,**H**) (*p < 0.05, **p < 0.01, ***p < 0.001, ****p < 0.0001). Data in scatterplots were assessed with linear regressions to generate the best-fit line. CL – central lateral, LP – lateral posterior, LD – lateral dorsal, DLG – dorsal lateral geniculate, PO – posterior, VPM – ventral posteromedial, VPL – ventral posterolateral, RT – reticular thalamus, VM – ventromedial. FGS – Facial grimace score.

For a more detailed analysis of thalamic stroke injury, thalamic nuclei were segmented from the chosen images using ABBA (Aligning Big Brains & Atlases) (Chiaruttini et al. 2024) with the Allen 3D Brain Atlas V3p1 (Wang et al. 2020), implemented in FIJI. Imported immunofluorescence images were assessed for the left/right and dorsal/ventral sectioning angle, positioned at the correct anterior-posterior location along the atlas, and then corresponding landmarks between the immunofluorescence images and atlas regions were manually selected using the Big Warp feature to register the 3D Allen Brain Atlas to brain sections. Overlaid ROIs generated from the registered atlas were imported into MATLAB for further analysis.

Allen Brain Atlas and stroke core ROI were imported into MATLAB for quantification. The extent of injury in each thalamic region was quantified as the ratio of the stroke core area within that nucleus to the total area of the nucleus, representing the percentage of the thalamic nucleus affected by the stroke core. Fluorescence intensity for each nucleus was calculated as the ratio of area-normalized fluorescence intensity in the right thalamic nucleus to that in the left thalamic nucleus.

### Statistical Analysis

Data are expressed as mean ± standard error of the mean (SEM) unless otherwise stated. All statistical tests were performed using GraphPad Prism version 10.3.1 (GraphPad Software, Boston, Massachusetts). Longitudinal behavioral analyses were performed using repeated measures Two-Way ANOVA with stroke status and time as the two independent variables. Positive ANOVA (α = 0.05) were assessed using Tukey’s multiple comparison test.

Histological comparisons for inflammation in the whole thalamus between sham and stroke mice, and for the ventricle ratio, were tested using unpaired two-sample t-tests. Thalamic nuclei-specific comparisons were made by first running a Two-Way ANOVA with thalamic nucleus and either stroke status (**Fig. 3D,H**) or stroke group (**Fig. 4E,F**) as the two independent variables. Positive ANOVAs (α = 0.05) were assessed using Tukey’s multiple comparison test. Linear regressions were performed separately on sham mice and stroke mice (**Fig. 3E,F**) to assess any relationship between the facial grimace score at four weeks post-stroke and the intensity of Iba1 staining in the VPL or VPM thalamus (calculated as the ratio of the average intensity in the ipsilateral nucleus/average intensity in the contralateral nucleus).

**Figure 4.**
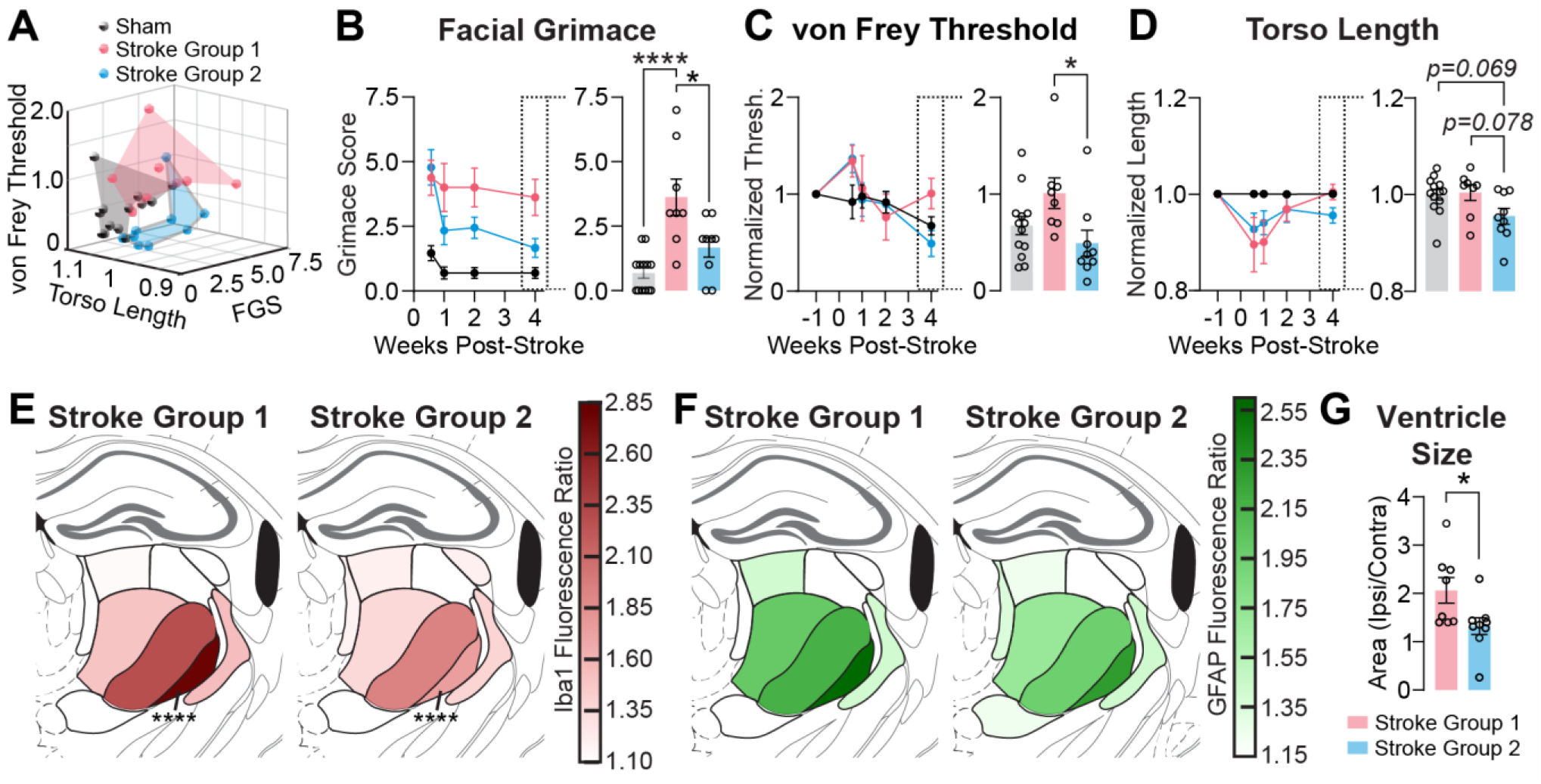
Mice with a thalamic photothrombotic stroke exhibit different behavioral phenotypes. **A)** Stroke mice fall into two distinct groups (red-SG1 and blue-SG2) based on facial grimace score, von Frey threshold, and torso length at four weeks post-stroke using k-means clustering performed on z-scores. Behavior scores come from Figure 2. Sham mice (not used for clustering) are superimposed (black). **B)** *Left:* Replotting grimace results from Figure 2A, after averaging separately mice in Stroke Group 1 (red), Stroke Group 2 (blue) and shams (black). *Right:* Differences in grimace scores at four weeks post-stroke between stroke groups and shams. **C)** *Left:* Replotting von Frey nocifensive behavior thresholds from Figure 2B, after averaging stroke groups separately. *Right:* von Frey thresholds at four weeks post-stroke between stroke groups and shams. **D)** *Left:* Replotting normalized torso lengths from Figure 2C, after averaging stroke groups separately. *Right:* Normalized torso lengths at four weeks post-stroke between stroke groups and shams. **E)** Mouse brain atlas showing the distribution of Iba1 staining quantified over different thalamic nuclei for stroke mice from Figure 3D, now separated across SG1 (left) and SG2 (right) according to the shown color map. Color values represent the ipsilateral/contralateral fluorescence ratio for each thalamic nucleus. **F)** Mouse brain atlas showing the distribution of GFAP staining quantified over different thalamic nuclei for stroke mice from Figure 3H, now separated across SG1 (left) and SG2 (right) according to the shown color map. Color values represent the ipsilateral/contralateral fluorescence ratio for each thalamic nucleus. **H)** The ipsilateral/contralateral ventricle size ratio from Figure 3J, separated by stroke group. Comparisons involving stroke groups and sham mice were compared with a One-Way ANOVA and Tukey’s multiple comparisons. Comparisons of fluorescence distributions across thalamic nuclei between SG1 and SG2 used a Two-Way ANOVA and Tukey’s multiple comparisons. Ventricle comparison between SG1 and SG2 used a two-sample t-test. *p <= 0.05, ****p < 0.0001. All other p values are specified.

Different stroke behavioral phenotypes were segmented by performing k-means clustering using the cityblock method in MATLAB on the facial grimace score, normalized torso length, and normalized von Frey pressure thresholds from only stroke mice, measured at four weeks post-stroke after calculating the z-score for each metric. Differences in these four-week behavior responses across sham, stroke group 1, and stroke group 2 were assessed with One-Way ANOVAs (α = 0.05), followed by Tukey’s multiple comparisons. An unpaired two-sample t-test was performed to identify difference in ventricle ratio between stroke groups.

For all statistical tests, a p-value <= 0.05 was considered statistically significant.

### Code Accessibility

Data and codes used to analyze the data are available upon request to the authors.

## Results

### Stroke Size Depends on Light Dose

We induced photothrombotic thalamic strokes in mice by inserting an optical fiber into the right VPL/VPM nuclei. We then injected the mice intraperitoneally with the photosensitive dye Rose Bengal (stroke group) or with saline (sham group). Laser light (532 nm) delivered through the optical fiber interacted with the dye producing oxygen free radicals, which trigger thrombosis and localized ischemia (Fluri, Schuhmann, and Kleinschnitz 2015).

To determine the optimal light dose to induce a VPL and VPM stroke, we exposed mice (n = 4–5 mice per group) to 95, 120, or 190 J/mm^2^ of 532 nm light (**Fig. 1A**). 2,3,5-Triphenyl-tetrazolium chloride (TTC) was used as an indicator of normoxia, staining oxygenated tissue red. It was applied to brain sections collected one day after light exposure. This revealed an unstained region in the thalamus of experimental mice, indicative of hypoxia caused by ischemic stroke. Mice in the sham group showed red staining throughout the brain, indicating normoxic tissue conditions in the thalamus and the absence of ischemia (**Fig. 1B**). The VPL and VPM localization of the stroke was confirmed by overlaying brightfield microscopy images with the Allen Brain Atlas using the ABBA tool in FIJI (**Fig. 1B, insets**). Increasing the duration of light exposure increased the volume of the stroke, which eventually engulfed most of the thalamus, as well as adjacent hippocampal, hypothalamic, and white matter tissue (**Fig. 1C**). Stroke volume was quantified, revealing a significant dose-dependent effect (F(2,19) = 6.442, p = 0.0073, Two-Way ANOVA) (**Fig. 1D**). No stroke was induced in sham mice at any light dose, indicating that light exposure alone does not cause ischemia (F(1,19) = 34.35, p < 0.0001, Two-Way ANOVA).

In subsequent experiments, we used a dose of 120 J/mm^2^ at 532 nm, which reliably produced a stroke in the VPL and VPM thalamic nuclei.

### Thalamic Stroke Results in Persistent Sensory Alterations

After optimizing the light dose, a separate group of mice (n = 13 sham, 17 stroke) was used to assess the longitudinal behavioral response to VPL and VPM thalamic stroke.

Assessing mice for the presence of spontaneous, rather than stimulus-induced pain is difficult, but recent studies indicate that the grimace scale provides an excellent measure. Here we measured the facial grimace score (Langford et al., 2010) at four days and one, two, and four weeks post-stroke. A significant effect of stroke (F(1,28) = 28.0, p = 9.5e-14, Two-ANOVA) and time (F(2.31, 62.62) = 8.94, p = 0.0002, Two-ANOVA) was observed (**Fig. 2A**). Most prominent grimace features were in the cheek and nose (**Fig. 2-1A–F**). At all time-points, mice with a thalamic stroke had a significantly higher grimace score than did sham mice (p < 0.0008, Tukey’s Multiple Comparisons).

We also assessed mechanical nociceptive thresholds using the von Frey filament test, which is one of the most commonly used methods in studies of pain and CPSP. We applied the von Frey filament to the plantar surface of the left hind paw, contralateral to the stroke. A significant effect of time post-stroke (F(3.31, 92.63) = 4.18, p = 0.0063, Two-Way ANOVA), but not stroke status was observed. At four days following the stroke, stroke mice displayed a significant hyposensitivity to stimulation, with a withdrawal threshold elevated by 35.5 ± 10.5% compared to sham mice (p = 0.044, Tukey’s Multiple Comparisons) (**Fig. 2B**).

To determine if thalamic strokes resulted in changes to spontaneous ambulation and posture, mice were placed in an open field setup. The distance traveled, time spent in the center of the open field, and maximum speed of movement (measurements usually used to quantify mobility, exploration, and anxiety behaviors) (Seibenhener and Wooten 2015) were not significantly different between shams and stroke mice (**Fig. 2-2 A–I**). However, mice with a stroke exhibited a hunched posture (F(1,28) = 6.697, p = 0.015, Two-Way ANOVA) (**Fig. 2C**) compared to shams, which has been used as an indicator of pain (Larson, Wilcox, and Fairbanks 2019; Sevcik et al. 2006), and was quantified here as a reduction in the torso length measured using Deeplabcut (Mathis et al. 2018), and normalized to pre-surgery length. This hunching behavior was significant up to one week following stroke induction, and most pronounced at four days post-stroke, with an 8.8 ± 3.1% reduction in torso length (p = 0.016, Tukey’s Multiple Comparisons). Despite displaying hunching, mice with a stroke did not perform worse on an accelerating rotarod test, which has been used to assess mobility, coordination, and stamina (Jaenisch et al. 2016) (**Fig. 2-2J–L**).

Overall, mice with VPL and VPM thalamic stroke exhibited a persistent facial grimace at four weeks post-stroke, a transient hyposensitivity to mechanical stimulation of the contralateral hind paw that resolved by one week post-stroke, and a reduced torso length that normalized by two weeks post-stroke. No correlation was observed between the results of multiple tests (Linear regressions: grimace vs. von Frey: p = 0.30, grimace vs. hunch: p = 0.45, hunch vs. von Frey: p =0.78) (data not shown) suggesting that performance on these behavioral tests were not related.

### Thalamic stroke results in sustained neuroinflammation, with microgliosis in the VPL thalamus correlating with grimace pain

To identify the extent of neuroinflammatory effects of the thalamic stroke, we used immunohistochemistry to label microglia (Iba1, ionized calcium-binding adaptor molecule 1) and reactive astrocytes (GFAP, Glial Fibrillary Acidic Protein). A typical staining pattern for a thalamic photothrombotic stroke includes an intense Iba1 signal at the center of the stroke core outlined by a glial scar high in GFAP signal (**Fig. 3A,B, Fig. 3-1A**). The coronal section at the center of the stroke core, chosen as the section that had the maximal cross-sectional core area and which was in the middle of the anterior-posterior range of sections with visible stroke core, was used to quantify the extent of neuroinflammatory staining induced in the thalamus of each. For sham mice, sections with residual inflammation from the insertion of the optical fiber (**Fig. 3A**, yellow arrow) were used as the center section. We used the ipsilateral/contralateral ratio of average fluorescence intensity to quantify inflammation in the stroke hemisphere, as we performed previously (Holden et al. 2021; Necula et al. 2021; F. S. Cho et al. 2022). Mice with a stroke had ~25% higher Iba1 staining in the ipsilateral thalamus than sham mice (t(28) = 3.73, p = 0.0009, unpaired t-test) (**Fig. 3C**). Segmentation of thalamic nuclei using ABBA showed that microgliosis was specifically localized to the VPL (p < 0.0001), VPM (p < 0.0001), and reticular (p = 0.005) thalamic nuclei relative to sham mice (Tukey’s Multiple Comparisons, **Fig. 3D**).

Microgliosis in the VPL was correlated with grimace scores at four weeks post-stroke (F(1,15) = 8.71, p = 0.01, linear Regression), a relationship not observed in sham mice (**Fig. 3E**). Notably, no relationship was observed between microgliosis in the VPM and grimace scores at four weeks post-stroke (**Fig. 3F**). We also assessed reactive astrocytes in the thalamus after stroke, and found ~35% higher GFAP staining across the ipsilateral thalamus in stroke mice compared to shams (t(28) = 5.64, p = 4.0e-6, unpaired t-test) (**Fig. 3G**). Nuclei-specific analysis revealed elevated GFAP staining in the VPL (p = 0.0001), VPM (p < 0.0001), posterior (p < 0.0001), and reticular (p = 0.01) thalamic nuclei (Tukey’s Multiple Comparisons) (**Fig. 3H**).

The lateral ventricles ipsilateral to optical fiber placement were enlarged in stroke mice compared to shams (t(28) = 2.86, p = 0.0079, unpaired t-test) (**Fig. 3I,J**), a clinical marker associated with increased risk and severity of neurological diseases (Ertekin et al. 2016; Zhou et al. 2015).

Stereotaxic coordinates and cross-sectional area of the center of the stroke core were measured (**Fig. 3-1C**). Mice with more ventral strokes had less sensitivity to von Frey filament stimulation of the left hind paw (F(1,15) = 6.46, p = 0.023, Linear Regression) (**Fig. 3-1D**), while more posterior strokes were associated with greater grimace scores (F(1,15) = 6.03, p = 0.027, linear Regression) (**Fig. 3-1E**), both at four weeks post-stroke. Grimace scores at four weeks post-stroke showed no correlation with inflammation in motor thalamic nuclei (**Fig. 3-1F–H**), indicating that the grimace response is not a result of motor impairment.

The stroke core (**Fig. 3B, Fig. 3-1B**), was used to quantify the physical extent of injury to the thalamus (**Fig. 3-1I**). The size of the stroke core was not related to grimace score at four weeks post-stroke (data not shown). Mice with more than 10% of the VPL thalamus hit by the stroke core exhibited significantly higher grimace scores at four weeks post-stroke compared to mice with less than 10% of the VPL thalamus injured (t(15) = 2.35, p = 0.033, unpaired t-test) (**Fig. 3-1J**), suggesting an estimate of the potential minimum lesion threshold in the VPL required to elicit spontaneous pain. The extent of stroke core injury in the VPM thalamus showed no correlation with grimace scores at four weeks post-stroke (data not shown).

### Persistent or resolving grimace pain at four weeks post-stroke is dependent on microgliosis in the VPL

Knowing that the clinical rate of CPSP after a thalamic stroke is ~50% (Treister et al. 2017), we aimed to determine 1) if our photothrombotic model also exhibits variability in pain responses, and 2) how behavioral differences among stroke mice correlate with neuroinflammation in the VPL and VPM.

We first applied a k-means clustering method to categorize stroke mice into two distinct behavioral populations—SG1 (stroke group 1, red) and SG2 (stroke group 2, blue) (**Fig. 4A**)—based on their grimace scores (**Fig. 2A**), von Frey thresholds (**Fig. 2B**), and torso length measurements (**Fig. 2C**) at four weeks post-stroke. We then replotted our longitudinal behavioral data (**Fig. 2**) separately for SG1 and SG2 mice (**Fig. 4B–D**), and found that there was a significant effect of stroke group at four weeks post-stroke (grimace: F(2,27) = 12.92, p = 0.0001, One-Way ANOVA—von Frey: F(2,27) = 3.83, p = 0.034, One-Way ANOVA—hunch: F(2,27) = 3.46, p = 0.046, One-Way ANOVA). Compared to SG2, mice in SG1 exhibited stronger facial grimace scores (p = 0.011, Tukey’s multiple comparisons test) (**Fig. 4B**) and greater hyposensitivity to paw stimulation (p = 0.029, Tukey’s multiple comparisons test) (**Fig. 4C**). Mice in SG2 had acute facial grimace that resolved (defined by no significant difference from shams) at four weeks post-stroke (**Fig. 4B**) and were more sensitive to paw stimulation than mice in SG1 (**Fig. 4C**). No significant difference was observed between SG1 and SG2 in torso length at four weeks post-stroke (p = 0.078, Tukey’s multiple comparisons test) (**Fig. 4D**).

We then investigated whether SG1 and SG2 also differed in stroke-induced neuroinflammation and injury to the thalamus. Mice in SG1 had 61% higher Iba1 staining in the VPL compared to those in SG2 (p < 0.0001, Tukey’s multiple comparisons test) (**Fig. 4E**), consistent with our earlier finding that microgliosis levels in the VPL correlate with persistent pain, as reflected by facial grimace scores (**Fig. 3E**). No significant differences were observed between SG1 and SG2 mice in Iba1 levels within other thalamic nuclei (**Fig. 4E**), in the amount of GFAP within thalamic nuclei (**Fig. 4F**), nor across the thalamus as a whole for Iba1 or GFAP (**Fig. 4-1A**). These findings identify microgliosis the VPL as the sole neuroinflammatory correlate of persistent grimace behavior.

The ipsilateral/contralateral ratio of lateral ventricle area was 56% greater in SG1 compared to SG2 mice (t(15) = 2.38, p = 0.03, unpaired t-test) (**Fig. 4G**), indicating more severe injury to the ipsilateral thalamus in SG1 animals. The cross-sectional area of the stroke core was not different across stroke groups (**Fig. 4-1B**), nor was there any difference in the amount of each thalamic nucleus affected by the core (**Fig. 4-1C**).

## Discussion

Thalamic stroke leads to the development of central post-stroke pain (CPSP) in approximately 50% of patients (Treister et al. 2017), profoundly reducing quality of life (Henry, Lalloo, and Yashpal 2008);however, how to prevent or reverse this pain remains unknown. While rodent models of CPSP after a thalamic hemorrhage exist (Hiraga et al. 2020; Gritsch et al. 2016), studies with these models have not confirmed the development of spontaneous pain, a key characteristic of CPSP in humans. Moreover, ischemic events account for approximately 87% of all strokes (Tsao et al. 2023); however, a preclinical model of CPSP induced by ischemic thalamic stroke has yet to be established. We filled this gap by developing a novel mouse model of CPSP using photothrombotic induction of thalamic stroke, and evaluated it for pain behaviors representative of the clinical condition.

### Relevance to clinical CPSP

The clinical presentation of CPSP is heterogeneous, often involving abnormalities in both spontaneous and evoked sensory experiences. Spontaneous pain, reported by approximately 85% of CPSP patients, is commonly described as aching, throbbing, stabbing, shooting, pricking, squeezing, freezing, burning, or lacerating (Klit, Finnerup, and Jensen 2009). Allodynia—the perception of non-noxious stimuli as painful—is present in ~90% of CPSP patients, most commonly in response to mechanical (~56%) and cold (~56%) stimuli (Paolucci et al. 2015). Sensory hyposensitivity is also common (Klit, Finnerup, and Jensen 2009), with ~50% of patients having tactile hypoesthesia (decreased touch sensitivity) (Greenspan et al. 2004). Pain can occur in various body regions, including the trunk, limbs, face, and affect one hemi-body or the entire body (Klit, Finnerup, and Jensen 2009).

We found that thalamic ischemic stroke in mice strongly elicited spontaneous pain, as evidenced by increased facial grimace scores (**Fig. 2A**), while responses to mechanical hind paw stimulation (**Fig. 2B**), posture (**Fig. 2C)**, and movement (**Fig. 2-2**) were variable. Mice in SG1 had spontaneous pain (persistent grimace) persisting to four weeks post-stroke, whereas mice in SG2 did not, relative to shams (**Fig. 4B**). We found that 47% (8/17) of mice fell within SG1, mimicking the clinical rate of CPSP following thalamic stroke (Treister et al. 2017). Because no persistent motor deficits were observed (**Fig. 2-2**), and the lack of correlation between inflammation in motor thalamic nuclei and grimace scores (**Fig. 3-1F–H**), we are confident that the observed grimace reflects pain rather than impaired facial motor control. Chronic facial grimace and hyposensitivity to mechanical stimuli correlated with more posterior and ventral strokes, respectively (**Fig. 3-1D,E**), characteristics which are also linked to a higher incidence of CPSP in humans (Treister et al. 2017; Krause et al. 2012). Mice with a greater grimace also showed greater microgliosis in the VPL thalamus (**Fig. 3E**), suggesting the role of chronic neuroinflammation in pain development, aligning with literature implicating purinergic microglial signaling in sustained post-stroke pain (Ma, Luo, and Wang 2022; Kuan et al. 2015). Lateral ventricle size, a clinical marker of increased risk and severity of neurological diseases (Ertekin et al. 2016), was also increased ipsilateral to stroke in mice with more spontaneous pain (**Fig. 4G**). Mice in SG2 tended to not strongly exhibit these clinically relevant histological markers (**Fig. 4E,G**), and their grimace resolved by four weeks post-stroke (**Fig. 4B**). Stroke size, which has not been related to clinical pain severity (G. Andersen et al. 1995), was also not related to pain (**Fig. 4-1B**), and no significant linear relationship existed between the stroke size and behavioral test performance (data not shown). Importantly, we found that the VPL was highly involved in the grimace pain phenotype whereas the VPM showed no such relationship (**Figs. 3E,F, 4E**).

One hypothesis for CPSP mechanisms implicates injury-induced loss of descending inhibition from the thalamus to the spinal cord ((Bud) Craig 1998). Central pain, including CPSP, has been treated with medial thalamotomy (Franzini et al. 2019) to remove the affective-motivational component of pain which may be hyperactive after injury-induced loss of descending inhibition to spinothalamic neurons from the sensory thalamus. The medial thalami in our mice were injury-free (**Fig. 3, Fig. 3-1I**), suggesting that grimace could be mediated through this region. The reticular thalamus (RT) provides GABAergic inhibition to the medial thalamus from neurons medially in the RT (Usrey and Sherman 2023; Clemente-Perez et al. 2017), a region with neuroinflammation in our stroke mice (**Fig. 3**). Stroke-induced neuroinflammation is sufficient to alter RT neuron excitability (Paz et al. 2010). Thus, a loss of RT-mediated inhibition of medial thalamic neurons represents an alternative mechanism for pain, warranting future electrophysiological investigation.

In summary, the photothrombotic model effectively recapitulates key features of CPSP after thalamic ischemic stroke, including heterogeneous persistent pain phenotypes, and offers a valuable tool for elucidating the mechanisms underlying CPSP.

### Expansion of the current CPSP toolbox

Standard preclinical CPSP studies use collagenase-induced thalamic hemorrhage (Hiraga et al. 2020; Gritsch et al. 2016). However, hemorrhage is more severe than ischemia (Chiu et al. 2010; K. K. Andersen et al. 2009), even though hemorrhagic stroke patients recover faster (Paolucci et al. 2003; Chae, Zorowitz, and Johnston 1996). This paradoxical observation suggests that stroke-induced cellular and circuit plasticity differs between hemorrhage and ischemia, and may not elicit CPSP through the same mechanisms. Our photothrombotic model of CPSP provides a platform to rigorously test these hypotheses.

Ischemic stroke has been previously applied to study CPSP (De Vloo et al. 2017). Arterial occlusions are most common, but are not thalamus-specific, with damage to cortical, thalamic, and sub-thalamic structures (Ueda et al. 2019). This extensive lesion volume is ill-suited for investigating thalamocortical circuits involved in CPSP (Gucer, Niedermeyer, and Long 1978; Vierck et al. 2013). Consequently, most prior ischemic stroke models of CPSP have not specifically assessed pain initiation within the thalamus.

The potent vasoconstrictor endothelin-1 can induce strokes of various size throughout the brain (Fluri, Schuhmann, and Kleinschnitz 2015)—an advantage this model shares with our photothrombotic model. Endothelin-1 has been suggested to act as a neurotransmitter (Fluri, Schuhmann, and Kleinschnitz 2015), and induces axonal sprouting and astrocytosis (Shihara et al. 1998), making it difficult to pinpoint whether electrophysiological results are from the ischemia or secondary endothelin-1 effects. While both photothrombosis and endothelin-1 serve as valuable research tools, the photothrombotic model offers greater precision for investigating thalamic circuit alterations implicated in CPSP.

### Limitations of the photothrombotic thalamic stroke model

Limitations of this model stem from the photothrombotic technique and clinical CPSP characteristics not recapitulated in the tested mice. Photothrombosis creates more edema than typically observed in clinical ischemic strokes (Sommer 2017), and the implications of this remain unclear (see below). In addition, the stroke penumbra from photothrombosis is very small (Sommer 2017), meaning the type and extent of neuroinflammation here may not fully mimic patients (implications below). We measured spontaneous pain in stroke mice (**Fig. 2A**), but did not observe the mechanical sensitivity changes that are a hallmark of hemorrhagic rodent and monkey CPSP models (Gritsch et al. 2016; Nagasaka et al. 2017), nor did we test hot or cold sensitivity. Therefore, as of yet, our model is not suitable to be used to understand the allodynia experienced by ~60% of CPSP patients (Ri 2022).

Some chronic thalamic neuroinflammation was induced in shams from optical fiber placement (**Fig. 3A,B**), although at a lower level than in stroke mice. Still, sham mice displayed lower von Frey thresholds at four weeks compared to baseline (p = 0.033), and it remains unclear whether this reflects a habituation effect or results from optical fiber placement. Regardless, ischemia in the thalamus has a much greater impact on persistent spontaneous pain emergence (**Fig. 2A**), acute hyposensitivity to mechanical stimulation of the contralateral hind paw (**Fig. 2B**), and acute hunched posture (**Fig. 2C**), not reproduced by the optical fiber alone.

### Future investigations

The development of this model of CPSP enables new, in-depth investigations into the cellular and circuit mechanisms of CPSP onset. Because the photothrombotic model failed to elicit mechanical hypersensitivity in the contralateral hind paw, a study directly comparing the pain emergence between collagenase-induced hemorrhage and photothrombosis-induced ischemia would be beneficial to identify model-specific neuroinflammation and circuit mechanisms responsible for the different types of pain. Brain edema following ischemic stroke is common, and a greater edema volume is related to worse neurological outcomes (Dostovic et al. 2016), but it is unclear how the increased edema from photothrombosis may result in the observed pain phenotypes. The small penumbra created by photothrombosis (Sommer 2017) could reduce the number of reversibly damaged neurons. More studies are needed to understand whether it is the reversibly damaged neurons within the penumbra, or the undamaged neurons that previously synapsed with now dead neurons, which are responsible for any altered excitability resulting in pain.

Overall, the preclinical model of CPSP presented here is a useful addition to the researcher’s toolbox for studying CPSP because of its ability to induce spontaneous pain out to four weeks post-stroke, and because of its ability to mimic predictive biomarkers of clinical CPSP. Future studies can apply this model to study mechanisms that govern pain emergence and chronification, and subsequently test novel therapeutics for preventing CPSP.

