## Supplemental Data for "Photothrombotic Ischemic Thalamic Stroke in Mice Recapitulates Spontaneous Pain Features of Central Post-Stroke Pain in Humans"

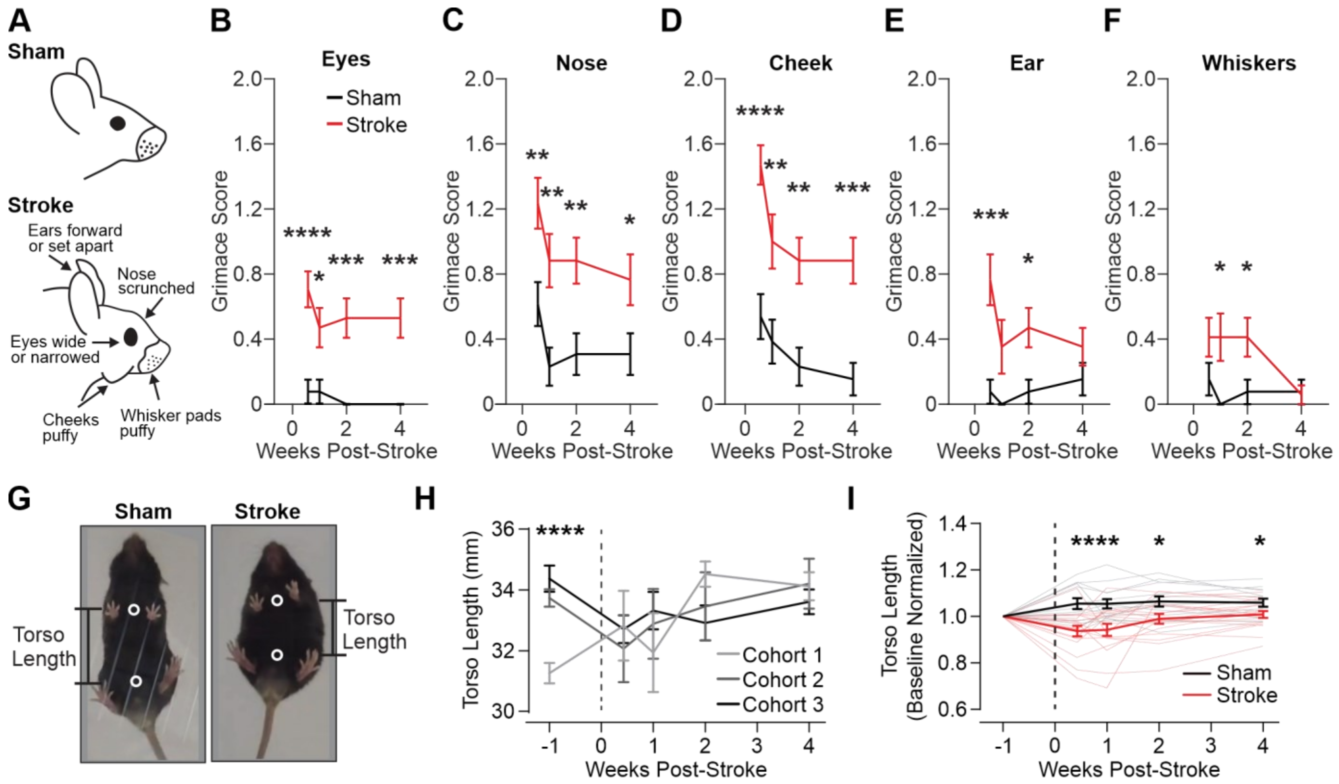


**Figure 2-1. Additional information about sensory tests performed**. **A)** An example drawing of grimace features (or lack thereof) exhibited by sham and stroke mice. **B**) Average eye grimace score for sham (black) and stroke (red) mice. **C**) Average nose grimace score for sham (black) and stroke (red) mice. **D**) Average cheek grimace score for sham (black) and stroke (red) mice. **E**) Average ear grimace score for sham (black) and stroke (red) mice**. F**) Average whisker grimace score for sham (black) and stroke (red) mice**. G)** Illustration examples of a sham, normal mouse and a mouse with a stroke that moves with a hunch. **H)** Torso length of mice across the three cohorts, with sham and stroke mice pooled together. It was identified that Cohort 1 was recorded at a lower camera resolution and frame rate, leading to different results and data normalization inaccuracies. Normalizing each cohort to the average of shams overcame this issue. **I)** Torso length normalized to each mouse's baseline torso length out to four-weeks post-stroke. This demonstrates that the reduced torso length from the Cohort 1 baseline artificially increases the average normalized torso length across all mice if within cohort normalization is not performed. Thick traces represent mean ± SEM for each group (stroke: red, sham: black). Data show all points from min to max, Mean ± SEM. For statistical comparisons, a Two-Way ANOVA ($\alpha = 0.05$) assessed differences across sham vs. stroke and over time, followed by Tukey's multiple comparisons performed on data after a positive ANOVA (*p < 0.05, **p < 0.01, ***p < 0.001, ***p < 0.0001).

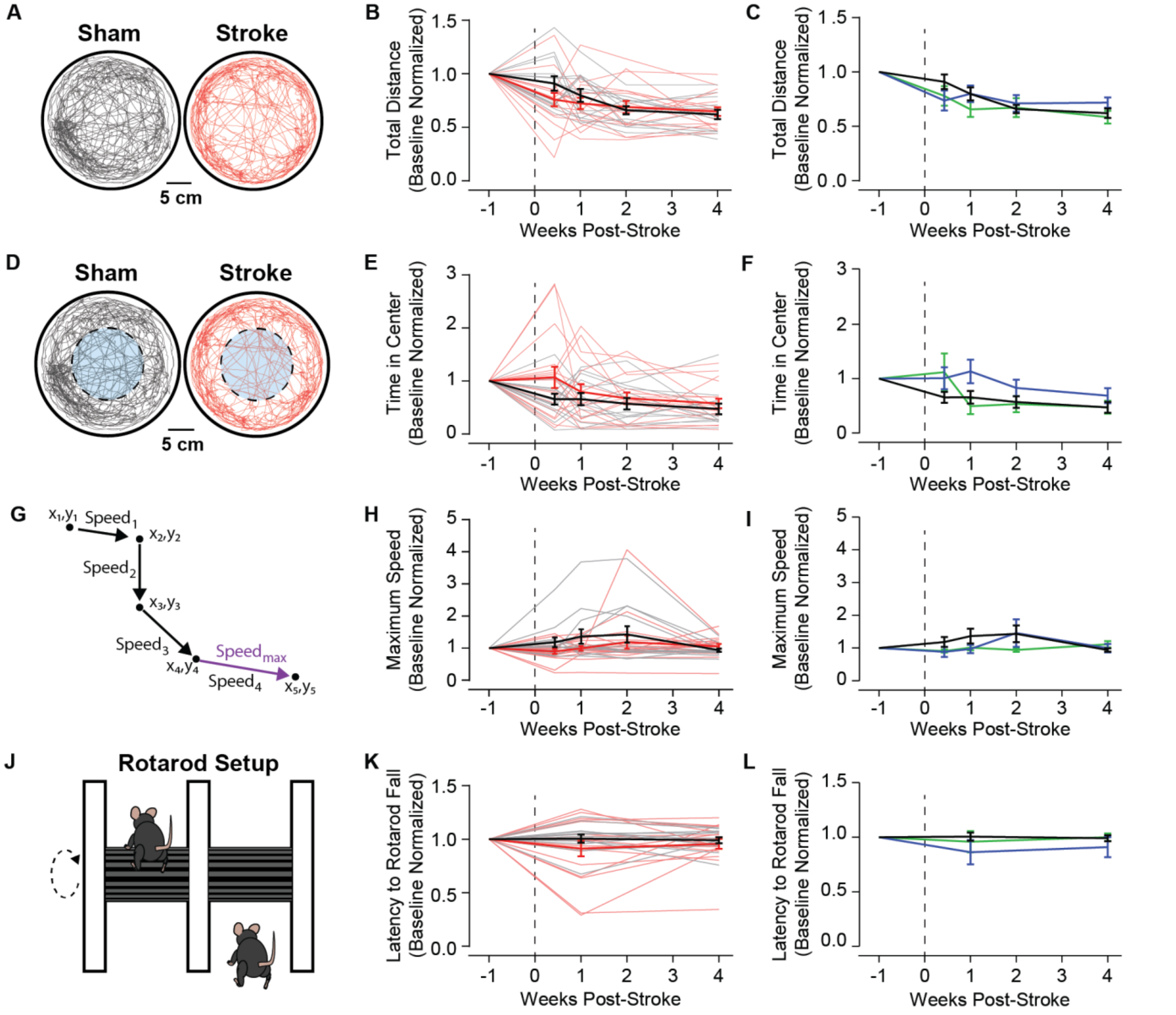


**Figure 2-2. Motor response to stroke**. **A)** The location of mice in an open field is tracked to calculate the total distance, a measure of total activity and movement capabilities, for sham (black) and stroke (red) mice. **B)** Total distance that mice traveled over 10 minutes in a cylindrical open field setup, normalized to baseline, shown for up to four-weeks post-stroke. **C)** Total distance results with mice separated into stroke groups identified in Figure 4. **D)** The location of sham (black) and stroke (red) mice are tracked and the amount of time spent within the center of the open field (dashed line) is quantified, where less time in the center is associated with greater anxiety. **E)** Time that mice spent in the center of the open field cylinder, normalized to baseline, shown for up to four-weeks post-stroke. **F)** Time in the center of the cylinder with mice separated into stroke groups identified in Figure 4. **G)** Maximum velocity across motion bouts is calculated for each video, providing a readout of any difficulty moving. **H)** The maximum velocity of mice in the open field, normalized to baseline, out to four-weeks post-stroke. **I)** Maximum velocity with mice separated into stroke groups identified in Figure 4. **J)** Rotarod setup where mice are placed on a rotating arm, and latency to falling off the rod is quantified. At least one hour after training the mice to stay on the rotating arm (3 trials at 8 rpm, up to 5 minutes), with testing performed on an accelerating rod (3 to 30 rpm, up to 5 min) for 3 trials. **K)** The average latency that it took mice to fall across rotarod testing trials, normalized to baseline, shown for up to four-weeks post-stroke. **L)** Latency to falling off the rotarod with mice separated into stroke groups identified in Figure 4.

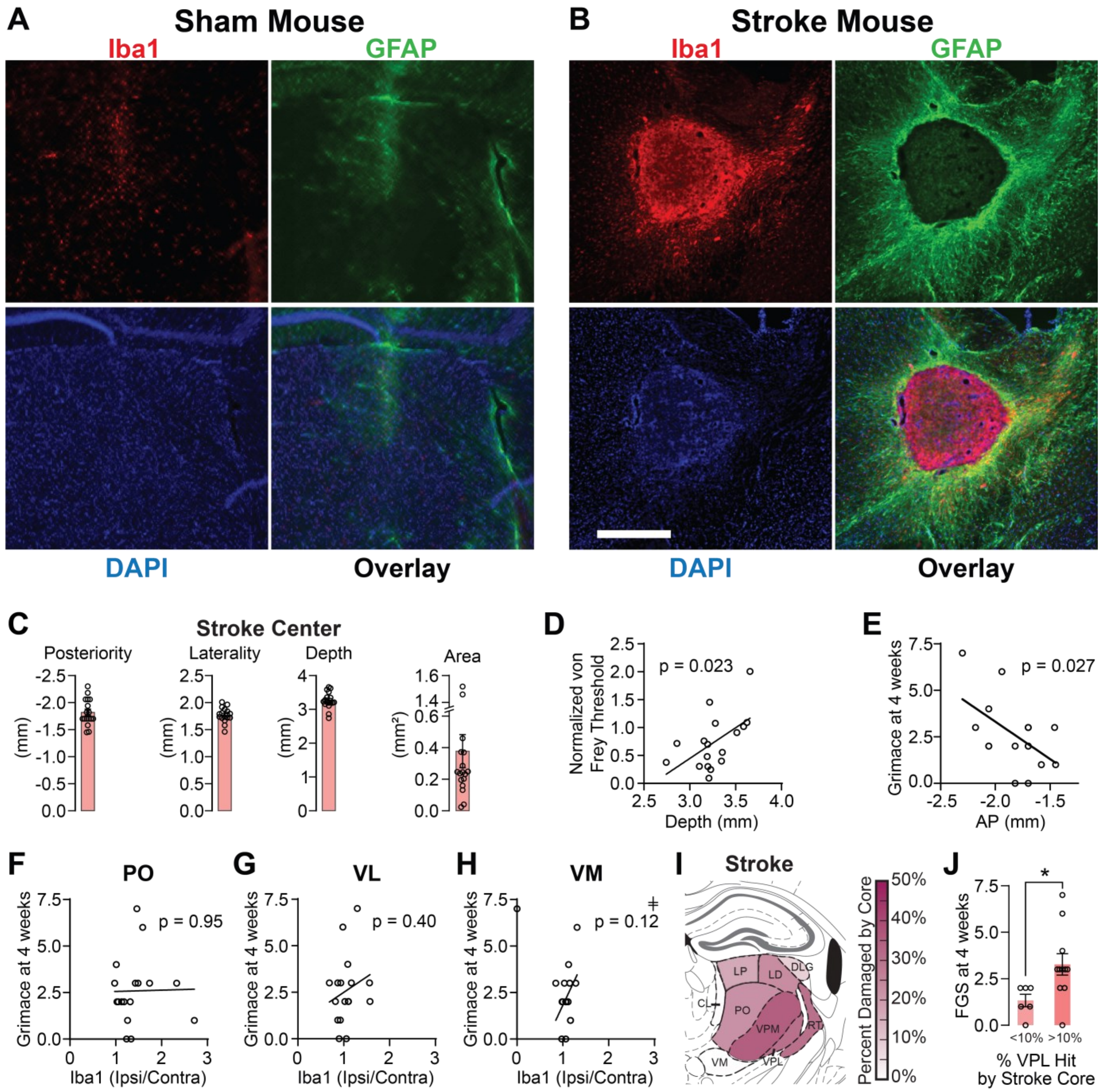


**Figure 3-1. Stroke cores are composed of a center high in microglia surrounded by astrocytic scarring**. **A)** Sham mouse center section from Figure 3, zoomed in and centered on thalamic inflammation from the optical fiber. Each channel is shown individually (Iba1 – red, GFAP – green, DAPI – blue) and overlaid. **B)** Stroke mouse center section from Figure 3, zoomed in and centered on the stroke core. Scale bar – 0.5 mm. **C)** Coordinates of the approximate center of the stroke and cross-sectional area of the stroke core for each mouse. **D)** Scatterplot of the normalized von Frey threshold of the contralateral hind-paw at four-weeks post-stroke versus the depth of the stroke core. A linear regression best-fit line is plotted with the regression p-value. **E)** Scatterplot of the facial grimace score at four-weeks post-stroke versus the anterior-posterior location of the stroke core. A linear regression best-fit line is plotted with the regression p-value. **F, G, H)** Scatterplots of the facial grimace scores at four-weeks post-stroke versus Iba1 fluorescence in the posterior (F), ventral lateral (G), and ventromedial (H) thalamic nuclei. Fluorescence is calculated for each nucleus as the ipsilateral/contralateral ratio. Linear regression best fit lines are shown with the p-values. **I)** Mouse brain atlas showing the distribution of the stroke core as the percent surface area of each thalamic nucleus that was hit according to the shown color map. **J)** Grimace scores of mice with 10% or more of the VPL thalamus hit by the stroke core are compared to grimace scores from mice with less than 10% of the VPL thalamus hit by

the core. Percent damage threshold was determined based on the threshold that maximally differentiated the facial grimace score at four-weeks post-stroke. Data represent all points from min to max, Mean ± SEM. Linear regressions were performed on scatterplot data. Two-group comparisons used an unpaired two-sample t-test. *p <= 0.05. All other p-values are specified. ǂIn one mouse, VM was not present in the analyzed section, resulting in a ratio of zero. This mouse was removed from the linear regression.

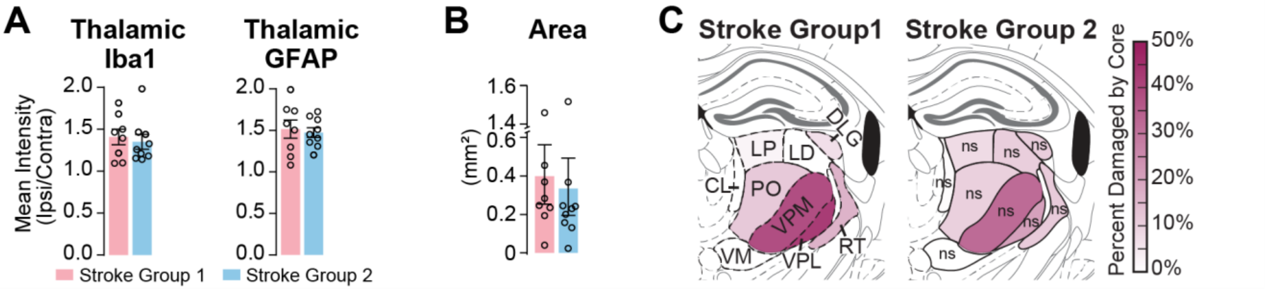


**Figure 4-1. Mice with a thalamic photothrombotic stroke exhibit different behavioral phenotypes. A)** Average fluorescence intensity of neuroinflammatory markers (ipsilateral/contralateral ratio) from Figure 3C,G, over the whole thalamus and separated across identified stroke groups from Figure 4. **B)** Cross-sectional area of the stroke core center from Figure 3-1C, separated by stroke group. **C)** Mouse brain atlas from Figure S3I showing the distribution of the stroke core as the percent surface area of each thalamic nucleus that was hit, now split across SG1 (*Left*) and SG2 (*Right*), according to the shown color map. Comparisons between SG1 and SG2 used two-sample t-tests. Nuclei-specific comparisons were performed using a Two-Way ANOVA between stroke group and thalamic nuclei. ns = no significance.
